# Cell type-specific biogenesis of novel vesicles containing viral products in human cytomegalovirus infection

**DOI:** 10.1101/2020.12.10.420711

**Authors:** Samina Momtaz, Belen Molina, Luwanika Mlera, Felicia Goodrum, Jean M. Wilson

**Affiliations:** Department of Cellular and Molecular Medicine, University of Arizona, Tucson, AZ 85721; Department of Immunobiology, University of Arizona, Tucson, AZ 85721; BIO5 Institute, University of Arizona, Tucson, AZ 85721

**Author notes:** **Co-Corresponding Authors:** Jean Wilson, PhD, Professor, Cellular and Molecular Medicine, Bio5 Institute, University of Arizona, 1501N Campbell Ave, Life Sciences North, Rm459, Tucson, AZ 85724, Felicia Goodrum, PhD, Professor, Immunobiology, BIO5 Institute, University of Arizona, 1657 E. Helen St., Thomas W. Keating Bldg, Rm 425, Tucson, AZ 85721.

## Abstract

Human cytomegalovirus (HCMV), while highly restricted for the human species, infects an unlimited array of cell types in the host. Patterns of infection are dictated by the cell type infected, but cell type-specific factors and how they impact tropism for specific cell types is poorly understood. Previous studies in primary endothelial cells showed that HCMV infection induces large multivesicular-like bodies that incorporate viral products including dense bodies and virions. Here we define the nature of these large vesicles using a recombinant virus where UL32, encoding the pp150 tegument protein, is fused in frame with green fluorescent protein (GFP, TB40/E-UL32-GFP). Cells were fixed and labeled with antibodies against subcellular compartment markers and imaged using confocal and super-resolution microscopy. In fibroblasts, UL32-GFP-positive vesicles were marked with classical markers of MVBs, including CD63 and lysobisphosphatidic acid (LBPA), both classical MVB markers, as well as the clathrin and LAMP1. Unexpectedly, UL32-GFP-positive vesicles in endothelial cells were not labeled by CD63, and LBPA was completely lost from infected cells. We defined these UL32-positive vesicles in endothelial cells using markers for the cis-Golgi (GM130), lysosome (LAMP1), and autophagy (LC3B). These findings suggest that virus-containing MVBs in fibroblasts are derived from the canonical endocytic pathway and takeover classical exosomal release pathway. Virus containing MVBs in HMVECs are derived from the early biosynthetic pathway and exploit a less characterized early Golgi-LAMP1-associated non-canonical secretory autophagy pathway. These results reveal striking cell-type specific membrane trafficking differences in host pathways that are exploited by HCMV.

**Importance:** Human cytomegalovirus (HCMV) is a herpesvirus that, like all herpesvirus, that establishes a life long infection. HCMV remains a significant cause of morbidity and mortality in the immunocompromised and HCMV seropositivity is associated with increased risk vascular disease. HCMV infects many cells in the human and the biology underlying the different patterns of infection in different cell types is poorly understood. Endothelial cells are important target of infection that contribute to hematogenous spread of the virus to tissues. Here we define striking differences in the biogenesis of large vesicles that incorporate virions in fibroblasts and endothelial cells. In fibroblasts, HCMV is incorporated into canonical MVBs derived from an endocytic pathway, whereas HCMV matures through vesicles derived from the biosynthetic pathway in endothelial cells. This work defines basic biological differences between these cell types that may impact the outcome of infection.

## Introduction

Human cytomegalovirus (HCMV) is a beta-herpesvirus that is characterized by its ability to establish a lifelong latent infection in humans with the potential for reactivation^1^. HCMV is prevalent worldwide, with a seroprevalence ranging from 45 to 100%, depending upon geographic location ^2^. In healthy individuals, HCMV infection is typically asymptomatic ^2, 3^. However, in immunocompromised individuals, such as stem cell or organ transplant recipients, HCMV reactivation or primary infection can result in high morbidity and mortality ^2, 4^. HCMV is vertically transmitted to developing fetuses, and approximately 1:150 children are born with congenital HCMV infection in the US ^5^, which can result in hearing impairment, microcephaly and neurodevelopmental delays ^6^. Asymptomatic seropositivity has also been linked to increased risk for age-related, chronic inflammatory pathologies, including vascular disease, frailty and immune dysfuntion ^7–11^. Currently, there is no vaccine for HCMV, and antivirals fail to eliminate the latent reservoir of virus because the drugs only target actively replicating virus ^12, 13^. Understanding HCMV biology and the mechanisms by which the virus replicates is important for developing strategies to control virus-related pathology and disease.

While HCMV is highly restricted in its tropism for the human species, a wide variety of cells are susceptible to infection within the human host, including fibroblasts, hematopoietic progenitor cells (HPCs), myeloid-lineage hematopoietic cells, smooth muscle cells, epithelial cells and endothelial cells (ECs). Fibroblasts and epithelial cells are major targets of the virus as these cells play a crucial role in the multiplication of the virus ^14^. Fibroblasts have been the primary model for studying HCMV replication because of the ability of these cells to support robust productive replication. However, HCMV establishes a chronic, low level persistence in endothelial cells ^15–17^. While productive HCMV infection in fibroblasts is well understood, we understand much less about the biology of infection in ECs. ECs are important targets of infection that undoubtedly contribute to HCMV dissemination and pathogenesis as ECs comprise the interface between the ciruclating blood and organs. Infection of the endothelium increases the recruitment and extravasation of monocytes and decreases vascular permeability ^18–23^. Further, proinflammatory signaling from the infected endothelium has been postulated to contribute to vascular disease ^24–26^.

In HCMV-infected human microvascular endothelial cells (HMVECs), we have observed the formation of large vesicles resembling multivesicular bodies (MVBs) that contain both virions and dense bodies (DBs) ^15^. The incorporation of HCMV products into these MVB-like vesicles suggests that they are a either a means of egress for the virus or a host response, resulting in a dead-end for infection if their contents are targeted for lysosomal destruction. Viruses containing disruptions in the HCMV gene, *UL135*, have defects in the incorporation of virions and DBs into the MVB-like vesicles, suggesting that pUL135 may direct incorporation of virus and DBs into these vesicles, possibly for egress. The existence of a viral gene that directs the incorporation of virus products into vesicles suggests that this may not represent an unfortunate dead-end cellular response to infection, but a direct goal.

In this study, we investigated the origin and identity of the MVB-like vesicles that incorporate viral products in ECs and fibroblasts. Using a virus where the pp150 tegument protein, encoded by *UL32*, is fused to the green fluorescent protein (UL32-GFP), we characterized the large vesicles that incorporate virus products containing UL32-GFP, such as virions and DBs. Interestingly, these large vesicles that contain virions in ECs do not have classical markers of MVBs, but do contain the lysosomal marker, LAMP1, and the *cis*-Golgi marker, GM130 on the limiting membrane, and the autophagy marker, LC3B, in the lumen of the vesicles. In contrast, in fibroblasts, UL32-GFP virions were incorporated into MVBs containing classical MVB markers. This unexpected result indicates that HCMV accesses distinct trafficking pathways in ECs and fibroblasts and suggests the possibility of distinct routes of egress in these cell types. Indeed, further characterization of these vesicles indicates that infection in ECs exploit early biogenesis/exocytic pathways, whereas infection in fibroblasts exploit endocytic pathways.

## Methods and materials

### Cells

Primary human lung microvascular lung endothelial cells (HMVEC) (purchased from Lonza, Walkersville, MD) were cultured in EGM-2 MV Bulletkit medium (Microvascular endothelial cell growth medium-2, Lonza) with 5% fetal bovine serum (FBS), 0.2mL hydrocortisone, 2mL hFGF, 0.5mL VEGF, 0.5mL R3-IGF-1, 0.5mL ascorbic acid, 0.5mL hEGF, and 100 U/mL penicillin. Human primary embryonic lung fibroblasts (MRC-5) (purchased from ATCC; Manassas, VA) were cultured in Dulbecco’s modified Eagle’s medium (DMEM) supplemented with 10% fetal bovine serum (FBS), 10mM HEPES, 1mM sodium pyruvate, 2mM L-alanyl glutamine, 0.1mM non-essential amino acids, 100 U/mL penicillin, and 100 μg/mL streptomycin. All cells were cultured at 37°C in 5% CO_2_.

### Viruses

Low passage strain of human cytomegalovirus TB40/E recombinant expressing UL32 fused to GFP, TB40/E-UL32GFP ^27^, was generous gifts from Dr. Christian Sinzger. Virus stocks were produced by the transfection of BAC DNA into fibroblasts and virus was isolated from cell-free culture supernants and concentrated through a sorbitol cushion. Virus was not serially propagated in fibroblasts. Virus titers were determined by TCID50 on fibroblasts as previously described^28^.

### Transmission Electron Microscopy

HMVEC or MRC-5 cells were mock-infected or infected at a multiplicity of infection (MOI) of 4 or 1, respectively, with centrifugal enhancement. Infection media was replaced 24 hours post infection for HMVECs and 6 hours post infection for MRC-5 with normal growth media for each respective cell type. All cells were harvested 5 days post infection (dpi) and fixed in 2.5% glutaraldehyde and 0.1M PIPES [piperazine-N, N′-bis (2-ethanesulfonic acid)] for 20 minutes. The fixed cell pellet was postfixed with osmium tetroxide in 0.1M PIPES and dehydrated in a graded series of alcohol. Pellets were infiltrated with resin and cut into 100 nm sections. The sections were floated on to copper grids and imaged using a Phillips CM-12s Transmission Electron Microscope. Cells were embedded and sectioned by the Arizona Research Laboratories, Arizona Health Sciences Center Core Facility.

### Immunofluorescence imaging

HMVEC and MRC-5 were seeded on to 12 mm glass cover slips in 24 well dishes one day prior to infection at a MOI of 4 or 1, respectively. Cells were processed for indirect immunofluorescence at 4 dpi for HMVECs and 2 dpi for MRC-5s. Cells were fixed in 4% paraformaldehyde in PBS (except for LC3B staining where we used ice-cold 100% methanol as a fixative) for 20 minutes. After washing with PBS, cells were incubated with 50mM ammonium chloride (NH_4_Cl) for 10 minutes to quench free aldehydes. Cells were blocked and permeabilized with 0.2% saponin in 10% FBS-containing 1XPBS for 30 minutes (Table 1). After blocking, the cells were incubated with primary antibodies for at least 2 hours. Primary antibodies were prepared by using antibody dilution buffer according to the manufacturer’s instructions. After rinsing with 1X PBS for at least three times, cells were incubated with secondary antibodies (Table 1) diluted in antibody dilution buffer, as per antibody datasheet, for 1 hour in a dark chamber. For methanol fixation of cells, we used anti-GFP secondary antibody since methanol quenches the fluorescence of GFP ^29^. Post staining, DNA was stained with DAPI according to manufacturer’s instructions (Molecular Probes). Coverslips were mounted using Prolong Diamond Antifade Mounting agent without DAPI (Invitrogen) according to manufacturer’s instructions. For Super resolution-structured illumination microscopy (SR-SIM) imaging, cells were cured for 60 hours in dark chamber before imaging. Confocal images were obtained with a Zeiss LSM880 inverted confocal microscope (Zeiss, Jena, Germany) with a 63X Plan Apo 1.4 NA oil immersion objective. All images were further processed using NIH-ImageJ ^30^. Representative single plane images with 0.5μm thickness were adjusted for brightness and contrast. Image galleries were created with Adobe Photoshop software (Adobe, San Jose, CA). SR-SIM images were obtained using a Zeiss ELYRA S.1(SR-SIM) Super Resolution microscope with a 63X Plan-Apochromat 1.4 NA objective. SIM processing and channel alignment were rendered using ZEN imaging software. Quantification of the vesicles was conducted using Image J. In brief, invert look-up tables (LUTs) of single plane images in two channels were processed and counted for coincidence of vesicles using point tools. At least one hundred vesicles were counted for each marker. Statistical analysis was performed using the Student’s t-test and error is indicated as SEM.

**Table 1:** Antibodies used in this study.

| Antibody | Species<sup>a</sup> | Source | Clone | Immunofluorescence |
| --- | --- | --- | --- | --- |
| CD63 | M | DSHB | H5C6 | 3µg/mL<sup>b</sup> |
| LBPA | M | Echelon | ML062915-21 | 1:100 |
| Clathrin | R | Cell signaling | 4796S | 1:50 |
| EEA1 | R | Cell signaling | 3288 | 1:100 |
| Rab5 | M | BD transduction | 610725 | 1:50 |
| ALIX | M | Invitrogen | MA1-83977 | 1:200 |
| Rab7 | R | Cell signaling | 9367 | 1:100 |
| P230<sup>c</sup> | M | BD transduction | 611280 | 1:400 |
| Rab6 | R | Cell signaling | 9625 | 1:400 |
| Rab11 | R | Cell signaling | 5589 | 1:100 |
| Rab14 | R | Sigma-Aldrich | R0656 | 1:200 |
| AP-2 | M | Abcam | ab2807 | 1:100 |
| AP-1<sup>d</sup> | M | Sigma-Aldrich | A4200 | 1:100 |
| AP-3 | M | DSHB | SA4 | 5µg/mL<sup>e</sup> |
| LAMP1 | R | Abcam | ab62562 | 1:500 |
| LC3B<sup>f</sup> | R | Cell signaling | 3868 | 1:200 |
| GM130 | M | BD transduction | 610822 | 1:100 |
| Anti-GFP | M | DSHB | G1-c 2ea | 5 µg/mL<sup>g</sup> |
| Alexa Fluor 647 | N/A | Thermo Fisher |  | 1:1000 |
| Alexa Fluor 568 | N/A | Thermo Fisher |  | 1:1000 |

<sup>a</sup> R, Rabbit; M, Mouse.
<sup>b</sup> original concentration 71µg/mL
<sup>c, d</sup> A generous gift from Dr. Samuel Campos, University of Arizona.
<sup>e</sup> original concentration 52 µg/mL
<sup>f</sup> fixed with ice-cold 100% methanol
<sup>g</sup> original concentration 239 µg/mL Infrared Imaging System (LI-COR, Lincoln, NE).

## Results

### UL32-GFP-positive vesicles in HMVECs are not classic MVBs

HCMV infection in endothelial cells (HMVECs) and fibroblasts induces the formation of large vesicles containing intraluminal vesicles that resemble multivesicular bodies (MVBs). By electron microscopy, we observed that virions and dense bodies are incorporated into these MVBs (Fig. 1). Previous studies have shown that MVB biogenesis and incorporation of viral products are controlled by different viral proteins in different cell types, such as pUL135 in endothelial cells and pUL71 in fibroblasts ^15, 31^. The requirement of viral proteins in the incorporation of virus into MVBs suggested that this is a virus-directed outcome. MVBs induced by infection and their biogenesis in the context of HCMV infection have not been characterized. To better define the MVBs induced by HCMV infection, we infected primary human microvascular endothelial cells (HMVECs) and MRC5 embryonic lung fibroblasts with a recombinant TB40/E strain engineered to express a variant of the UL32/pp150 tegument protein that has been fused to green fluorescent protein (GFP) ^32^. Multiplicities of infection (MOI) were chosen to infect ~50-60% of the cells for each cell type. Due to the different kinetics of infection in HMVECs and fibroblasts, cells were fixed and stained for confocal microscopy at 2 days post infection (dpi) and 4 dpi for fibroblasts and HMVECs, respectively. These time points were chosen such that infection would be at a similar same stage in each cell type; early in the late phase of infection where the viral assembly compartment (VAC) was apparent, but prior to severe cytopathic effect.

**Fig. 1:**
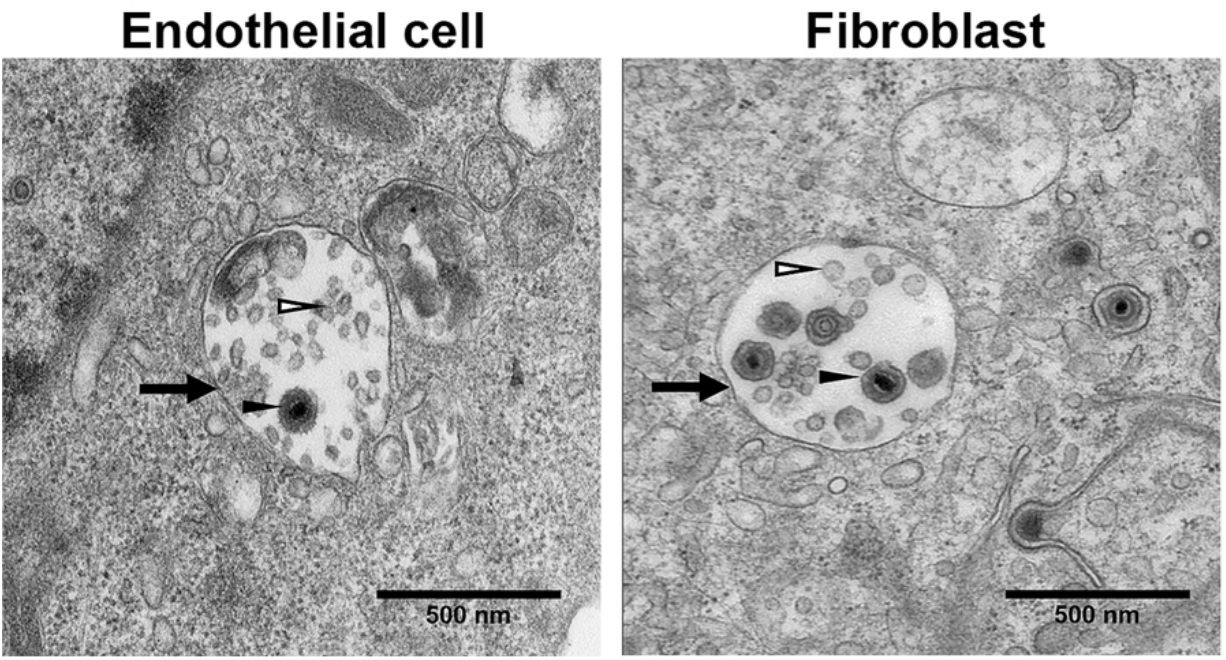
Virions are incorporated into MVBs. TB40/E-infected HMVECs (MOI, 4) or fibroblasts (MOI,1) were imaged by transmission electron microscopy. Multi-vesicular bodies (black arrows) are present in the cytoplasm. The lumen of the MVBs contain virions (filled arrowheads) and ILVs (open arrowheads). Scale bars, 500nm.

To define the composition of MVBs that incorporate viral cargo, we labeled cells with the classic MVB markers CD63 and LBPA. CD63 is a Tetraspanin-group protein that is present in MVBs or late endosomes (LEs) and at the cell surface ^33^.

Lysobisphosphatidic acid (LBPA) is present on the membrane of the intraluminal vesicles (ILVs) and is used as marker of ILVs in MVBs ^34^. In uninfected HMVECs (Fig. 2A and C, top panels) and fibroblasts (Fig. 2A and C, bottom panels), CD63 and LBPA are distributed on punctate perinuclear structures. In infected HMVECs, neither CD63 (Fig. 2A, top) nor LBPA (Fig. 2C, top) co-localized with UL32-GFP. Strikingly, LBPA was undetectable in infected HMVECs, although adjacent uninfected cells contain LBPA (Fig. 2C). In contrast, in infected fibroblasts, both CD63 (Fig. 2A, bottom) and LBPA (Fig. 2C, bottom) colocalized to large UL32-GFP-positive vesicles. Quantification of the vesicles in fibroblasts shows that 90% and 97% of UL32-GFP with colocalized with CD63 and LBPA, respectively compared to no colocalization of UL32-GFP with either marker in HMVECs (Fig. 2B and D). UL32-GFP labels the membrane of CD63-positive vesicles in fibroblasts, and also accumulated in the lumen of vesicles, hence likely represents the incorporation of UL32-containing virions and dense bodies (Fig. 2E) observed by electron microscopy (Fig.1). These results suggest that, while UL32-GFP containing vesicles in fibroblasts are classical MVBs, vesicles in HMVECs are atypical or non-classical.

**Fig. 2.**
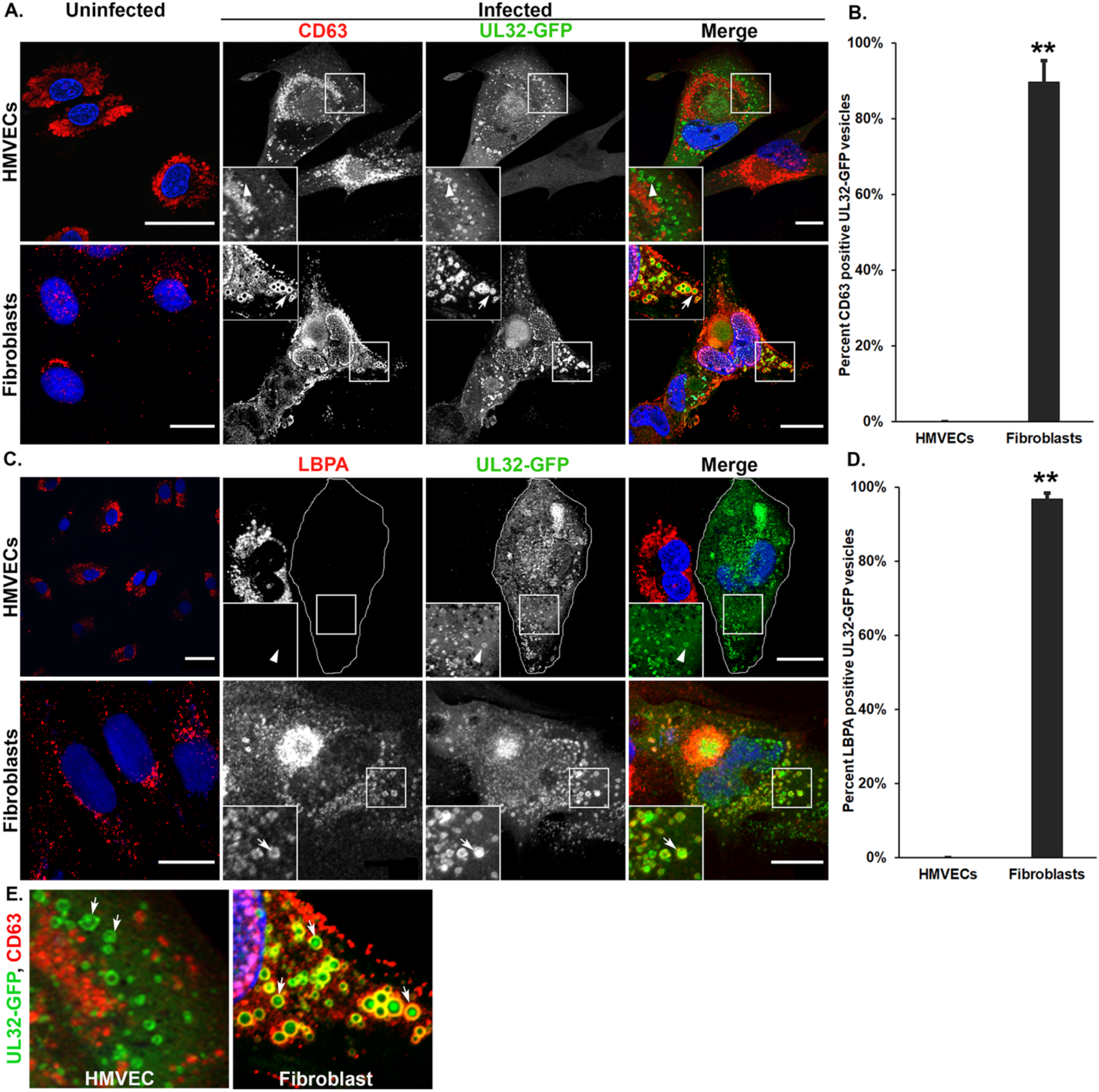
CD63 and LBPA show differential association with UL32-GFP-positive vesicles in HMVECs and fibroblasts. Uninfected or TB40/E-UL32-GFP-infected HMVECs (MOI, 4) or fibroblasts (MOI,1) were fixed at 96 and 48 hpi, respectively. (A) Cells were labelled with mouse anti-CD63 (red) and imaged by confocal microscopy. Nuclei are stained with DAPI. The large UL32-GFP-positive vesicles do not colocalize with CD63 in HMVECs (arrowheads, insets) but do colocalize with CD63 in fibroblasts (arrows, insets). (B) Quantification of the percentage of vesicles that contain the CD63 that are also positive for UL32-GFP, ** p< 0.01. (C) Uninfected and infected cells were labeled with anti-LBPA (red) and imaged by confocal microscopy. Infection results in loss of LBPA from infected HMVECs, and the infected HMVEC is outlined due to lack of LBPA staining. UL32-GFP positive vesicles are indicated by the arrowhead (inset). In fibroblasts, UL32-GFP positive vesicles colocalize with LBPA (arrows). (D) Quantification of the percentage of vesicles that contain the LBPA that are also positive for UL32-GFP, **p< 0.01. For each quantification, 700 to 900 vesicles were counted for each marker. E. UL32-GFP accumulates in the lumen of the large vesicles in both HMVEC and fibroblasts (arrows). Scale bars, 20μm.

### Clathrin heavy chain associates with UL32-GFP-positive vesicles

Clathrin acts as an integral component of both endocytic and biosynthetic cargo trafficking ^35,36^. Clathrin domains on MVBs receive Hrs-sorted ubiquitinated cargo for transfer to ESCRT-I and incorporation into the MVB ^37–39^. Previous studies have shown accumulation of clathrin near the VAC in infected fibroblasts, and this accumulation was decreased by inhibition of endocytosis ^40^. However, the localization of clathrin to MVB-like vesicles in the context of HCMV infection has not been examined. In uninfected HMVECs, clathrin is widely distributed on punctate cytoplasmic structures and is tightly localized to the perinuclear region in fibroblasts (Fig. 3). However, in infected HMVECs (Fig.3, top panels) or fibroblasts (Fig.3, bottom panels), clathrin is localized on the large (average size of 0.6-1.0μm) UL32-GFP-positive peripheral vesicles, (Fig. 3A, insets). Perinuclear clathrin accumulation in infected fibroblasts is consistent with previous findings that clathrin is re-localized to the VAC during infection ^40, 41^. However, the re-localization of clathrin to the VAC in infected HMVECs is less prominent. Quantification of the vesicles showed 81% of UL32-GFP-positive vesicles in infected HMVECs and 99% of UL32-GFP-positive vesicles in infected fibroblasts contained clathrin (Fig. 3B). These results demonstrate the accumulation of UL32-GFP on and within large, clathrin-positive, peripheral vesicles in both fibroblasts and HMVECs.

**Fig. 3.**
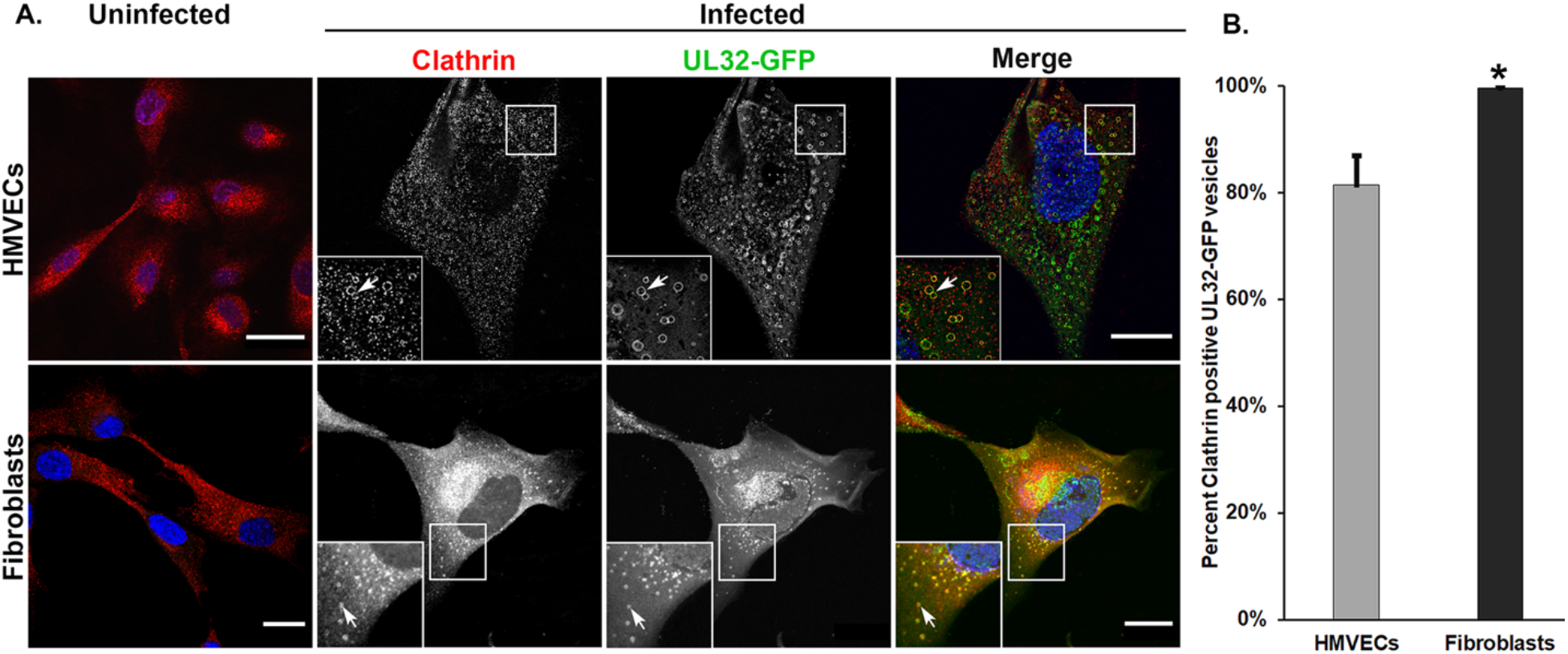
Clathrin heavy chain colocalizes with UL32-GFP-positive vesicles in HMVECs and fibroblasts. (A). Uninfected or TB40/E-UL32-GFP-infected HMVECs or fibroblasts labelled with anti-clathrin heavy chain (red). In uninfected cells, clathrin is distributed in diffuse puncta throughout the cell. There is substantial colocalization of clathrin heavy chain and UL-32-GFP vesicles (arrows, insets) in both HMVECs and fibroblasts. In fibroblasts, clathrin also accumulates in the VAC (lower middle panel). (B). Quantification of clathrin-positive, UL32-GFP positive vesicles. Thirteen hundred-vesicles were counted. *p<0.05. Scale bars, 20μm.

### UL32-GFP-positive vesicles lack common endosomal and biosynthetic compartment markers in HMVECs

Our results show that the UL32-GFP-positive vesicles in HMVECs, but not fibroblasts, lack the classical MVB markers, CD63 and LBPA. These findings suggest that UL32-positive vesicles in fibroblasts derive from the conventional endocytic pathway whereas the vesicles in HMVECs may originate from a distinct pathway. To further investigate this, we examined the localization of early and late endosomal, and biosynthetic compartment markers (Fig. 4).

**Fig. 4.**
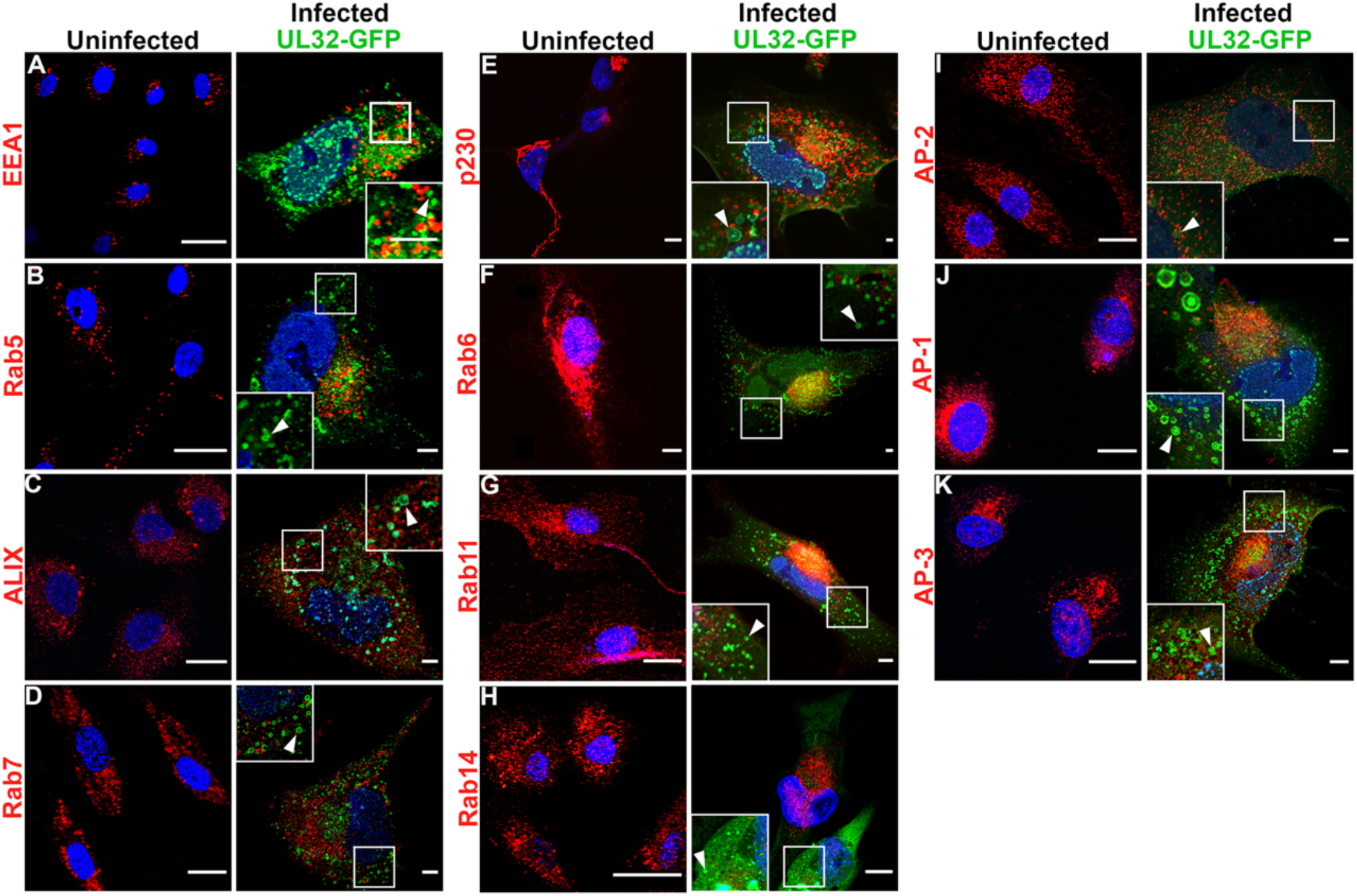
UL32-GFP-positive vesicles do not contain typical endocytic and biosynthetic trafficking markers in HMVECs. (A-K). Uninfected or TB40/E-UL32-GFP-infected HMVECs. Cells were labelled with (A) anti-EEA1 (early endosomes), (B) anti-Rab5 (early endosomes), (C) anti-ALIX (late endosomes), (D) anti-Rab7 (late endosomes, (E) anti-p230 (trans-Golgi), (F) anti-Rab6 (trans-Golgi), (G) anti-Rab11 (recycling endosomes), (H) anti-Rab14 (endosomes), (I) anti-AP-2 (plasma membrane), (J) anti-AP-1 (trans-Golgi), and (K) anti-AP-3 (trans-Golgi) antibodies and secondary antibodies conjugated to Alexa-fluor 647. The UL-32-GFP-positive vesicles (arrows) do not colocalize with any of these markers. Nuclei, blue. Scale bars, 20μm.

The early endosome (EE) is often marked by the presence of early endosomal antigen (EEA) 1 and the small GTPase Rab5, which mediates early endosome docking and fusion in association with phosphatidylinositol 3-phosphate (PI_3_P) ^42^. Previous studies have shown that HCMV infection in fibroblasts and HMVECs redistributes EEA1 to the VAC, suggesting a possible role of EEA1 in viral maturation ^43, 44^. In uninfected HMVECs, EEA1 and Rab5 are distributed on puncta in the cytoplasm (Fig. 4A-B), and neither colocalized with the UL32-GFP vesicles in infected HMVECs (Fig. 4B). These results indicate that UL32-GFP-positive vesicles in HMVECs do not derive from or maintain markers of the early endosomal pathway.

ALG-2 interacting protein-X (ALIX) binds LBPA, and CHMP4B, a component of the ESCRT-III complex, which is involved in membrane curvature and fission^45^. A previous study established that down-regulation of ALIX reduces the LBPA labeling by 50% ^46^. As LBPA was undetectable in infected HMVECs, we asked if ALIX localization is affected in infected HMVECs. In uninfected HMVECs, ALIX is localized on small cytoplasmic granules (Fig. 4C). In infected HMVECs, ALIX localization was unaltered, and did not colocalize with UL32-GFP vesicles (arrowhead, inset). We further tested for colocalization of the late endosome-associated small GTPase, Rab7. Rab7 regulates MVBs biogenesis and trafficking from early endosomes to late endosomes and lysosomes^47^. Uninfected HMVECs showed Rab7 labeling distributed throughout the cytoplasm and we observed no colocalization of Rab7 with UL32-GFP vesicles (Fig. 4D). In sum, these results indicate that none of the EE or LE associated markers colocalize with UL32-GFP-positive vesicles in HMVECs and suggest that these vesicles do not derive from the endosomal pathway.

Based on these results, we next tested if the UL32-GFP-positive vesicles in HMVECs could be derived from the biosynthetic pathway. We labeled infected HMVECs with the trans-Golgi marker, p230. A tubular network of p230 was observed in the perinuclear region of uninfected HMVECs, but this TGN marker did not colocalize with UL32-GFP-positive vesicles (Fig. 4E), suggesting that the UL32-GFP-positive vesicles do not derive from the trans-Golgi network.

Next, we tested the small GTPase, Rab6, which regulates retrograde transport from the endosomal compartment via the trans-Golgi to the endoplasmic reticulum^48^. A previous study found that Rab6 recruits UL32 to the viral assembly compartment by binding to dynein, a microtubule motor protein ^49^. Rab6 was scattered through the cytoplasm in uninfected cells and did not colocalize with UL32-GFP vesicles in infected HMVECs (Fig. 4F). This data suggests that endosomal and trans-Golgi membrane traffic is not involved in the biogenesis of these vesicles in HMVECs.

A previous study showed that HCMV assembly compartment formation alters the recycling endosomal Rab cascade marked by the small GTPase Rab11 ^50^. Rab11 was distributed as small punctate structures in the cytoplasm of uninfected cells and did not colocalize with UL32-GFP-positive vesicles in infected HMVECs (Fig. 4G). The small GTPase, Rab14, which is involved in the biosynthetic trafficking between Golgi and endosomes and the plasma membrane ^51^, also did not colocalize with UL32-GFP-positive vesicles in HMVECs (Fig. 4H). These data suggest that UL32-positive vesicles are not derived from endosomal recycling compartments.

Due to our observation of the presence of clathrin on the membrane of UL32-GFP-positive vesicles in HMVECs, we next asked if UL32-GFP positive vesicles contained clathrin-associated adaptor proteins: AP-2, AP-1, or AP-3. AP-2 binds to the phosphatidylinositol 2-phosphate (PIP_2_) in the plasma membrane and the cargo in clathrin-mediated endocytosis ^35^. AP-2 labeled small puncta in uninfected HMVECs, and this distribution did not change with infection in infected HMVECs (Fig. 4I). However, AP-2 was somewhat less intense and more diffuse, possibly due to infection-induced increase in cell size. AP-2 did not colocalize with UL32-GFP vesicles (Fig. 4I). AP-1 recruits clathrin to the TGN and contributes to the biogenesis of vesicles from the TGN ^52^. Uninfected HMVECs showed the small punctate structure of AP-1 in the perinuclear region (Fig. 4J), similar in appearance to p230 (see Fig. 4E). AP-1, like AP-2, also did not colocalize with UL32-GFP vesicles (Fig. 4J). AP-3, mediates the transport from TGN to LE or lysosome/Lysosome Related Organelles ^53^. As with all other adaptor proteins, AP-3 also did not colocalize with UL32-GFP-positive vesicles in infected HMVECs (Fig. 4K). Together, these findings demonstrate a striking lack of colocalization of UL32-GFP with common endosomal and biosynthetic markers and support the idea that these vesicles in HMVECs originate from a non-classical membrane trafficking pathway.

### UL32-GFP-positive vesicles associate with lysosomal, autophagy and early biosynthetic (cis-Golgi) markers in HMVECs

MVB-associated cargoes have two primary fates, i) fusion with the lysosomal compartment for the degradation, or ii) transport to the plasma membrane for exosomal release ^15, 54^. Our results show that UL32-GFP containing vesicles colocalized with classical MVB markers in fibroblasts but not in infected HMVECs. To further characterize these vesicles, we asked if UL32-GFP vesicles colocalize with the lysosomal compartment marker, LAMP1. Uninfected HMVECs showed elongated vesicular labeling of LAMP1 throughout the cytoplasm (Fig. 5A). In infected HMVECs, LAMP1 labeling was not substantially altered. However, the UL32-GFP containing vesicles colocalized with LAMP1 (Fig. 5A).

**Fig. 5.**
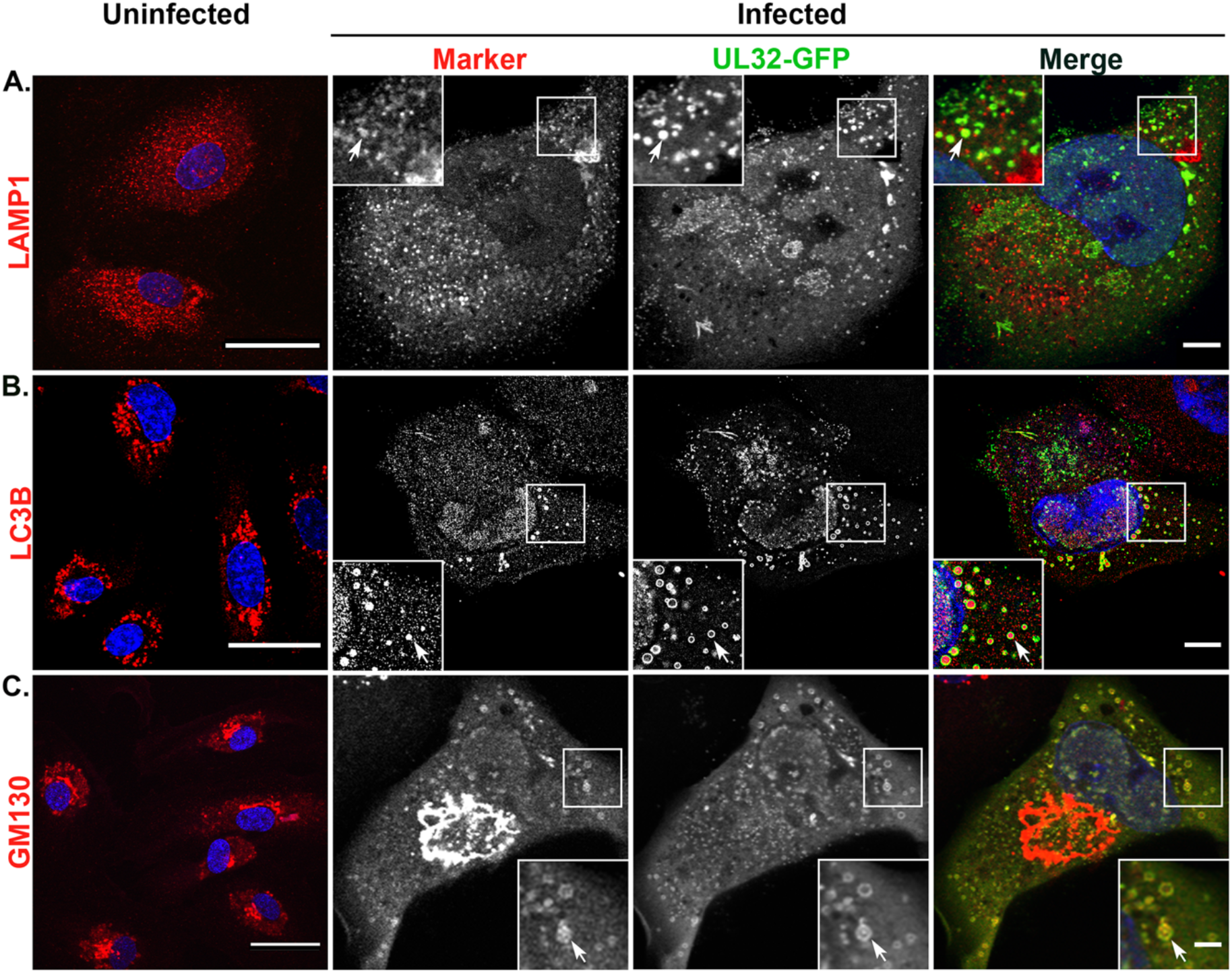
LAMP1, GM130, and LC3B localize to UL32-GFP-positive vesicles in HMVECs. Uninfected or TB40/E-UL32-GFP-infected HMVECs were labelled with (A) anti-LAMP1 (late endosomes-lysosomes), (B) anti-LC3B (autophagosomes), and (C) anti-GM130 (cis-Golgi) antibodies and secondary antibodies conjugated to Alexa-fluor 647 (red). Nuclei, blue. In uninfected cells, LAMP1 and LC3B are distributed throughout the cytoplasm. GM130 is localized to the perinuclear region, In infected cells, LAMP1, LC3B, and GM130 all colocalize on UL32-GFP-positive vesicles (arrows). Scale bars, 20μm.

A recent study reported that HCMV hijacks the autophagic component LC3B for envelopment of infectious virus particles, and knockdown of LC3B by shRNA demonstrated reduced viral production ^55^. Next, we asked if those non-classical MVBs in infected HMVECs contained LC3B. In uninfected HMVECs, LC3B was distributed in spherical cytoplasmic structures (Fig. 5B). In infected HMVECs, we observed LC3B in the lumen of the UL32-GFP vesicles (Fig. 5B). Luminal localization of LC3B within non-classical MVBs marked by UL32-GFP in HMVECs suggests that the biogenesis of MVBs in HMVECs may follow a non-classical secretory autophagy pathway, like LC3-dependent EV loading and secretion (LDELS).

Lysosomal storage vesicles have been described that are labeled by LAMP1, clathrin, and the cis-Golgi marker GM130, but are negative for LE markers ^56^. These vesicles sometimes also contain LC3B^57^. To determine if the large vesicles observed in HMVECs could be related to these structures, we next examined if the UL32-GFP vesicles contained GM130. In infected HMVECs, GM130 labeling was detected as a well-defined and characteristic ring-structure around the VAC (Fig. 5C). Further, GM130 colocalizes with the UL32-GFP vesicles in HMVECs (Fig. 5C). These finding indicate that the UL32-GFP vesicles may derive from an early biosynthetic, Golgi-mediated pathway.

### UL32-GFP containing vesicles are marked by LAMP1, but not LC3B and GM130 in fibroblasts

We next sought to analyze the association of LAMP1, LC3B, and GM130 with UL32-GFP-positive vesicles in infected fibroblasts. LAMP1 labelling in uninfected fibroblasts appeared as elongated vesicular structures in the cytoplasm (Fig. 6A). Infected fibroblasts demonstrated a scattered and fine granule labelling of LAMP1 around the VAC, consistent with previous observations (Fig. 6A) ^43^. However, UL32-GFP containing vesicles also colocalize with LAMP1 (Fig. 6A). LAMP1 has been reported to localize in the late endosomes (LEs), apart from their primary localization at the lysosomal compartments ^58–60^. This finding indicates that colocalization of UL32-GFP-positive vesicles with LAMP1 in infected fibroblasts may derive from either LEs or from the lysosomes.

**Fig. 6.**
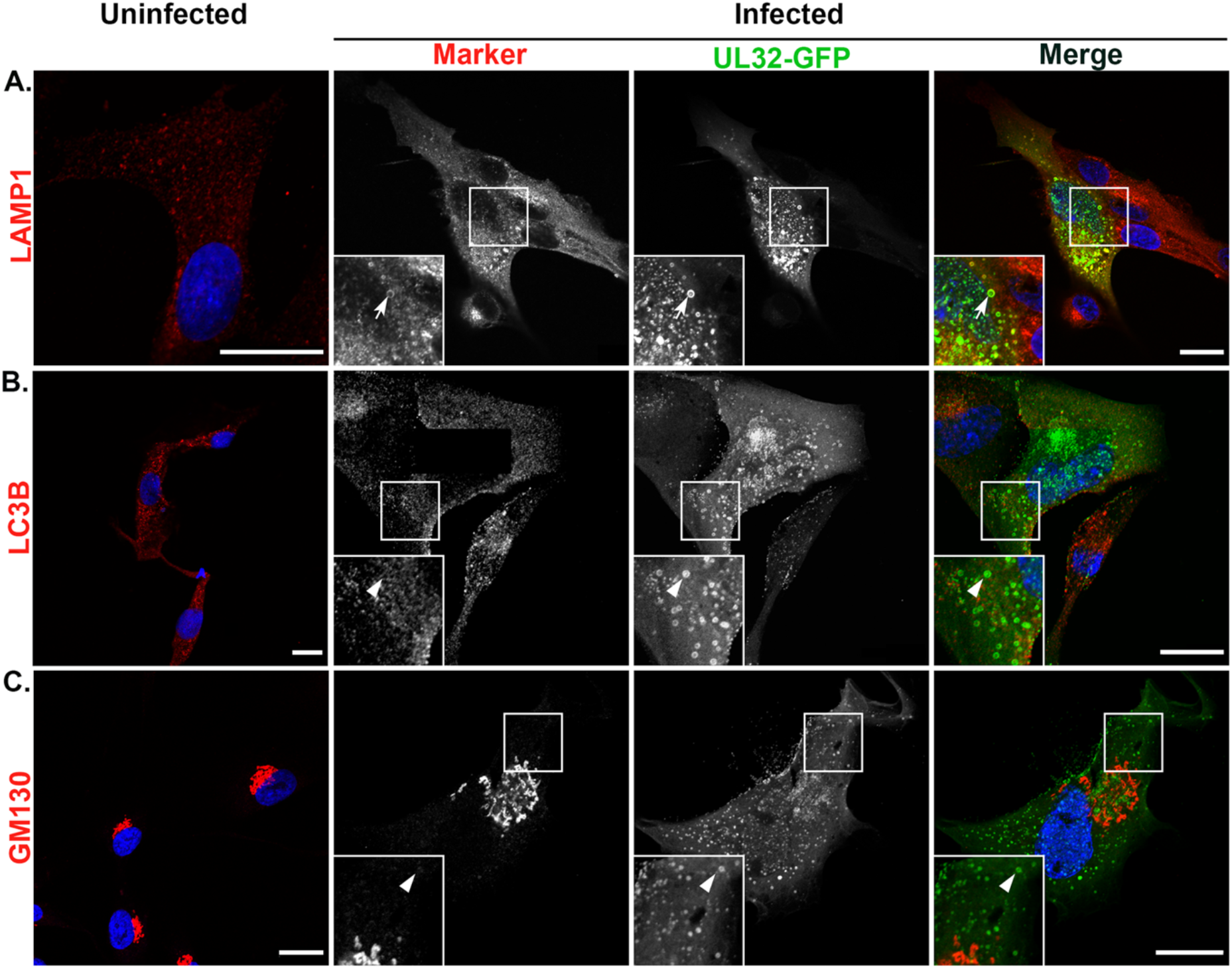
LAMP1, but not GM130 or LC3B, colocalize with UL32-GFP-positive vesicles in fibroblasts. Uninfected or TB40/E-UL32-GFP-infected fibroblasts were labelled with (A) anti-LAMP1 (late endosomes-lysosomes), (B) anti-LC3B (autophagosomes), (C) anti-GM130 (cis-Golgi) antibodies and secondary antibodies conjugated to Alexa-fluor 647 (red). Nuclei, blue. In uninfected cells, LAMP1 and LC3B are distributed in the cytoplasm. GM130 localizes to the perinuclear region. In infected cells, UL32-GFP vesicles colocalize with LAMP1 (arrows, A), but not with GM130 or LC3B (arrowheads, B, C). Scale bars, 20μm.

LC3B was present on vesicle structures throughout the cytoplasm in uninfected fibroblasts (Fig. 6B). However, LC3B localized predominantly to the VAC and no colocalization was detected with UL32-GFP vesicles in infected fibroblasts (Fig. 6B). In uninfected fibroblasts, GM130 localized in a tubular, perinuclear region typical of the cis-medial Golgi (Fig. 6C). In infected fibroblasts, GM130 was localized to the classic ring-like structure of the VAC and showed no colocalization with UL32-GFP vesicles (Fig. 6C).

We quantified the colocalization of LAMP1, LC3B and GM130 with UL32-GFP in both HMVECs and fibroblasts (Fig. 7). In HMVEC and fibroblasts, 92% and 99% of the UL32-GFP vesicles colocalized with LAMP1, respectively (Fig. 7A). However, LC3B and GM130 showed strong discordance between HMVECs and fibroblasts; 87 % of UL32-GFP-positive vesicles colocalized with LC3B in infected HMVECs, compared to none in infected fibroblasts (Fig. 7B). Similarly, 88% of UL32-GFP vesicles localized with GM130 in HMVECs, compared to none in infected fibroblasts (Fig. 7C). The absence of classic MVB markers on UL32-GFP vesicles in HCMV-infected HMVECs coupled with the presence of LC3B and GM130 suggest that these vesicles are derived from a distinct pathway from that associated with infection in fibroblasts.

**Fig. 7.**
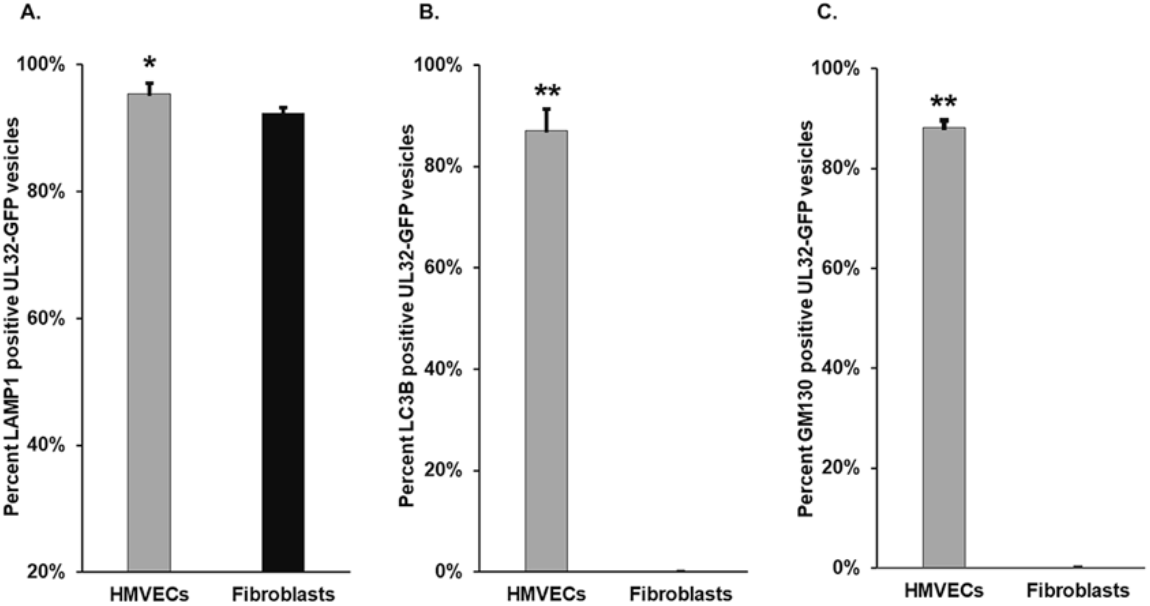
Quantification of LAMP1, LC3B, and GM130 positive UL32-GFP vesicles. The percentage of LAMP1 (A), LC3B (B), and GM130 (C) positive UL32-GFP vesicles on both cell types were illustrated. Two hundred to four hundred vesicles were counted for each marker. Most vesicles in both cell types contain LAMP1. However, fibroblasts do not contain LC3B or GM130. *p <0.05, **p< 0.01

## Discussion

HCMV establishes distinct patterns of infection depending upon the cell type infected. HCMV infection in fibroblasts results in a robust productive infection while vascular endothelium supports a smoldering, chronic infection that serves as a gateway of hematogenous dissemination to distant organs ^14, 61, 62^. However, the cellular mechanisms that distinguish these patterns and regulate post-entry tropism remain unclear. Defining the molecular and cellular basis for these distinct patterns of replication is fundamental to understanding the biology of infection in specific cell types and supports efforts to identify antiviral targets for controlling hematogenous HCMV spread. We previously observed the incorporation of virus products of replication, virions and dense bodies, in multivesicular-like bodies in endothelial cells ^15^. In this study, we used a virus in which the viral protein pp150 has been fused in frame with green fluorescent protein to define the membrane-bound vesicles incorporating virions and dense bodies. Remarkably, we find that endothelial cells exploit a specialized trafficking pathway that is associated with a LAMP1-mediated biosynthetic and atypical secretory autophagy trafficking, whereas fibroblasts route viral components and progeny through membranes that resemble classical MVBs.

We find that the membrane markers present on MVBs in infected fibroblasts represent classical MVBs that include CD63, LBPA, and LAMP1 ^33, 34, 54, 63–67^. It has been well established that other viruses, such as hepatitis C and hepatitis A use the ESCRT component Hrs, as well as VPS4 protein (AAA ATPase) and ALIX for incorporation into MVBs for export ^68, 69^. The presence of these markers on virus-containing MVBs in fibroblasts suggests that these vesicles originate from the canonical endocytic trafficking pathway. In the canonical endocytic pathway, the early endosome (EE) is the sorting station for cargo and determines its fate ^70^. EE mature to form late endosome (LE) by replacing EE resident Rab5 to Rab7 ^70^. Also during maturation, LEs incorporate endosomal membrane to generate ILVs containing MVBs ^33, 70^. Depending on the cargo, MVBs can then transport cargo either to the degradative lysosomes or for fusion with the plasma membrane for release of contents from the cell ^33, 54, 70^.

Unexpectedly, the virus-containing MVBs in HMVECs lack classical MVB markers, indicating their biogenesis from a pathway distinct from endocytic trafficking pathways. The loss of LBPA in the infected HMVECs also implies the generation of altered ILVs or altered lipid metabolism. This may be due to viral interference with LBPA synthesizing enzymes or their precursors, e.g. phosphatidyl glycerol ^68^. Altered ILV generation is supported by the lack of colocalization of these vesicles with the exosomal marker ALIX. A previous report showed ALIX-independent ILVs with altered biogenesis and shape, thus affecting the vesicle release pathway^46^. ALIX-independent generation of ILVs has also been observed with other viruses, e.g., Herpes simplex virus (HSV-1) and HIV^71, 72^. Moreover, loss of LBPA results in reduced retention of cholesterol ^73^, that blocks classical exosomal release^74^. Furthermore, other lipids, such as ceramides and their metabolite sphingosine-1 phosphate (S1P) have been implicated in MVB release ^68^. Thus, in microvascular endothelial cells, HCMV likely uses a distinct mechanism for egress, although a definitive role for these vesicles in egress remains to be determined. In cells from individuals with lysosomal storage disorders, membrane vesicles marked by clathrin, LAMP1, GM130, and LC3B exist that are not directed to lysosomal compartments ^57^. These vesicles are thought to be exaggerations of normal membrane trafficking pathways. The colocalization of LAMP1, GM130 and the cluster of LC3B in the lumen of the UL32-GFP vesicles supports the idea the vesicles induced by HCMV infection in endothelial cells are similar and are not targeted for degradation; hence they could be directed to fuse with the plasma membrane for release. This possibility remains to be tested.

Viral hijacking of degradative autophagy is commonly observed for enhanced cell survival, viral persistence, or for viral reactivation and release ^75^. HCMV in fibroblasts also hijacks autophagy during viral maturation ^55^. However, secretory autophagy has been adopted by several viruses, including poliovirus, human rhinovirus 2, coxsackievirus, zika virus, and Epstein Barr virus (EBV), as a route to exit host cells ^76–80^. These viruses utilize lipidated LC3B-rich membrane bound vesicles, derived from the autophagosome, to transport virus particles out of the cells ^76, 81^. Intriguingly, luminal localization of LC3B within the vesicles of HMVECs suggests a secretory autophagy pathway. While little is known about luminal LC3B dependent secretory autophagy, LC3-dependent EV loading and secretion (LDELS) has been reported ^82^. Interestingly, these vesicles accumulate ceramide, not LBPA, to facilitate membrane scission in an ESCRT-independent manner ^82^, which supports our finding of loss of LBPA in virus-containing vesicles. Most recently, SARS-CoV-2, has been shown to egress using a lysosomal pathway where the virus likely traffics to lysosomes from the Golgi/trans Golgi network or the ER/ER-Golgi intermediate compartment (ERGIC) ^83^.This egress pathway requires deacidification of the lysosomes so that the incorporated virus is not destroyed. HMVECs may use a similar egress pathway. However, the luminal LC3B in HCMV-containing vesicles also suggests the association with secretory autophagy, distinct from SARS-CoV-2 egress.

Based on our findings, we propose a model in which HCMV controls maturation and egress from distinct host pathways in infected HMVECs and fibroblasts (Fig. 8). The vesicles in infected fibroblasts contain CD63, LBPA, LAMP1, and clathrin, thus may originate from canonical endocytic trafficking pathways and represent the classical exosomal release pathway ^84^, resulting in productive infection. In contrast, vesicles in HMVECs contain clathrin, GM130, LAMP1, and luminal LC3B. This suggests that these vesicles derive from the early biosynthetic route. Luminal LC3B may promote virion incorporation and recruit ceramide to facilitate membrane scission and release as EVs. Our results suggest that HCMV has evolved to differentially utilize two distinct vesicle trafficking pathways in fibroblasts and endothelial cells.

**Fig. 8.**
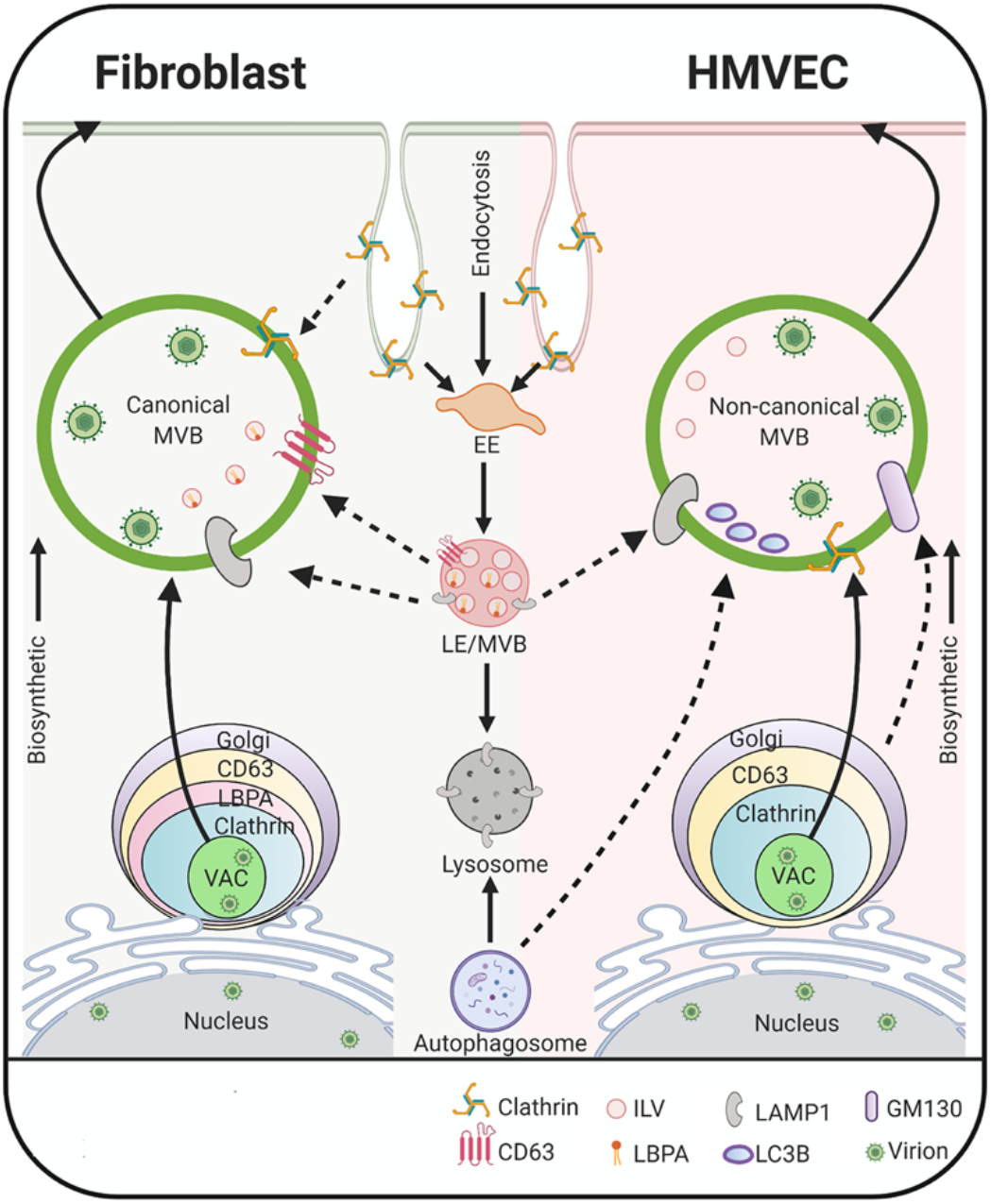
Proposed model. In fibroblasts, HCMV infection induces classical MVBs, suggesting that the virus exploits membrane trafficking in the endocytic pathway to promote viral incorporation into these vesicles, possibly for release via the exosomal pathway. In endothelial cells, HCMV infection generates MVBs that contain non-canonical markers, suggesting that the MVBs in these cells originate from the early biosynthetic pathway and viral infection expands the non-canonical secretory autophagy pathway. The solid arrows denote the known cellular pathways; dotted arrows denote pathways expanded by HCMV infection. The image was created with BioRender.com

## Acknowledgements

We acknowledge Dr. Christian Sinzger (Ulm University, Germany) for the kind gift of the UL32-GFP virus. We acknowledge Patricia Jansma, William Day and Douglas Cromey of the University of Arizona Imaging Cores for assistance with confocal and transmission electron microscopy. This work was funded by a National Institutes of Health/National Institute of Allergy and Infectious Disease grant AI131598 to FG and JMW.

